# The search for surrogacy in patient derived xenograft mouse trials: glass is less than half full

**DOI:** 10.1101/2020.12.03.409730

**Authors:** Hitesh B. Mistry

## Abstract

Despite the efforts of many within the drug development and clinical community surrogate biomarkers for patient survival have remained elusive in Oncology. This failure in part is attributable to there being a paucity of clinical trials showing a treatment effect on patient survival. Given this issue an alternative system to explore the surrogacy potential of biomarkers are large preclinical xenograft studies i.e. panel of patient derived xenografts or mouse clinical trials. In this study we explored the surrogacy potential of tumour burden biomarkers, current size of tumour and how its changed, preclinically in a large patient derived xenograft database which contains a diverse number of drugs/treatments (n=61) and xenografts (n=245). We found that of the possible 1830 two-arm mouse trials, 1103 showed a treatment effect on the preclinical end-point, disease progression, (p<0.05). Of these only in 30% did tumour burden markers fully capture the treatment effect on disease progression times i.e. satisfied a key condition for surrogacy. These results highlight that preclinically it is very challenging to find a surrogate marker based purely on measures of tumour burden.

## Introduction

The holy grail for mathematical/statistical modelling groups within Oncology drug development is predicting the clinical effectiveness of a new drug/treatment as early as possible. Clinical effectiveness here would be showing an improvement in patient survival of the new drug/treatment versus the standard of care.^1^ In order to make such predictions the community requires a biomarker which fully captures the treatment effect on patient survival, a surrogate biomarker.^2^ (Note, this biomarker need not be a single variable it could be a combination of numerous variables.) In order to find such a biomarker, we need clinical studies which show a difference in patient survival between treatments. These studies though are quite rare.^3^ Therefore, generating enough evidence clinically to have faith in the surrogate biomarker is a difficult task.

An alternative strategy to assess a potential surrogate biomarker could be to use preclinical xenograft studies. There are numerous advantages to exploring potential surrogate biomarkers preclinically. A key one being that numerous drugs could be tested at a fraction of the cost compared to the clinic. Currently, the industry is using large patient derived xenograft collections to explore the effectiveness of new treatments in what are termed as mouse clinical trials.^4,5^ These databases now also contain numerous drugs and so could be used to explore the potential of surrogate biomarkers. Indeed, one such study by Novartis^4^ contained 60 treatments and 100s of genetically diverse xenografts has been made available publicly. That study followed mice for almost a year until the disease had progressed and so contains rich time-series with an end-point. In this article we shall use this database to explore the surrogacy potential of biomarkers that capture the dynamics of tumour burden.

## Methods

Tumour volume time-series data was extracted from Gao et al.^4^ The volumes were then converted to diameters, assuming the tumours were spherical. In order to make the analysis clinically relevant the clinical, RECIST^6^, definition of target lesion progression, 20% increase over the minimum with a minimum increase of 5mm, was translated to the preclinical setting in the following way. The 20% increase over the minimum value seen was kept. The minimum increase value was set to the preclinical limit of quantification value of 1mm. Disease progression times (PFS) were then calculated as the time from treatment initiation to disease progression. Xenografts which had not progressed by the end of the observation period were considered right-censored.

1830 two arm trials were created by exploring all head-to-head drug possibilities. All trials were then assessed for a difference in treatment effect on PFS via the log-rank test. Only studies which had a p-value<0.05 were used for the biomarker analysis, described below.

Given that the xenograft is simply a single lesion the time-varying biomarkers considered are the current diameter value at time t, L(t), and how that value is changing over time, DLDT(t), defined as the difference in two consecutive measurements divided by the time difference between them. In order to assess whether these two biomarkers fully capture the treatment effect on PFS the following regression analysis framework was used.

First a Cox model without treatment arm as a covariate was fit to the data,

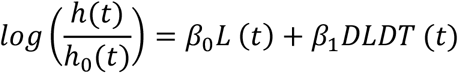

and then one with treatment arm,

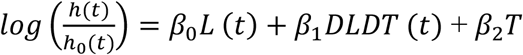

where *h*_0_(*t*) is the baseline hazard function, *L*(*t*), *DLDT(t)* and *T* represent the diameter, rate of change of the diameter and the treatment arm, respectively at time *t. β*_0_,*β*_1_ and *β*_2_ are regression coefficients to be determined. Note that in this model, since the covariates vary over time, the ratio *h*(*t*)/*h*_0_(*t*) is also time-dependent and the model is no longer a proportional hazards model. The likelihood ratio-test is then used to assess whether including treatment arm as a covariate improves model fit. If the p-value<0.05 then it was concluded that the biomarkers did not fully capture the treatment effect on PFS.

All analyses were done in R v3.4.0. Final data-sets and R scripts can be found here: https://github.com/HiteshBMistry/PreclinicalSurrogacyProject.

## Results

Of the 1830 trials created from the database 1103 showed a difference in PFS that gave a log-rank test p-value <0.05. A volcano plot showing how the p-value relates to the PFS HR can be seen in Figure 1. The studies plotted in green are the ones that showed a difference in PFS between the two treatment arms and those in red did not. The plot shows that the HRs of the studies being taken forward into the surrogate biomarker analysis cover a wide range.

**Figure 1:**
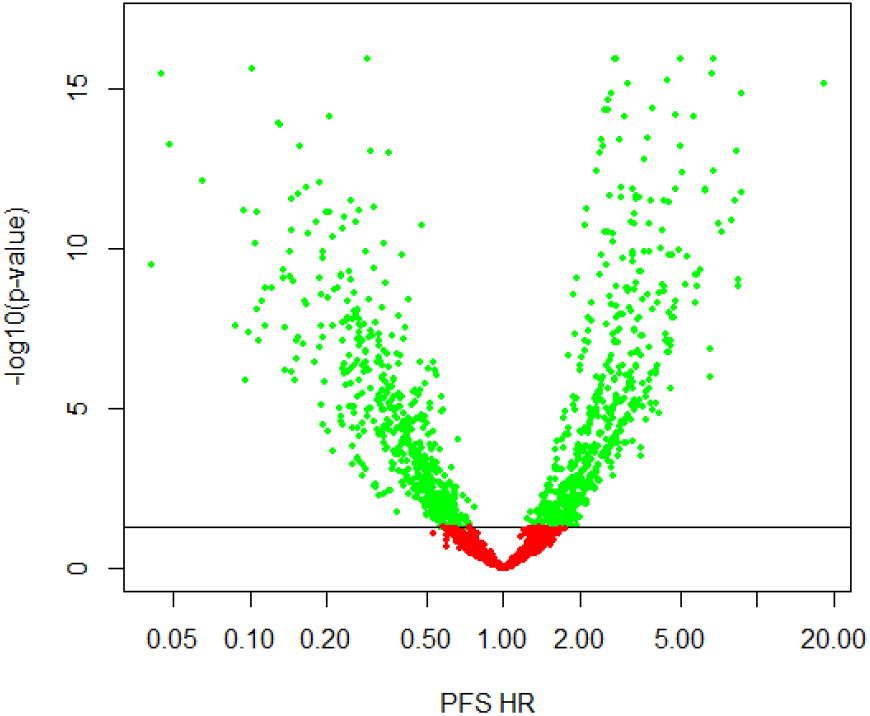
Plot showing the p-value and HR for each of the 1830 two-arm trials created. Those in green had a p-value<0.05 and those in red did not.

**Table 1.** shows the total number of mice, the number progressed and the median PFS time for each drug/treatment.

| **Treatment** | **Total N**<br>**(No. Progressed)** | **PFS (Days)**<br>**Median (95% CI)** |
| --- | --- | --- |
| LEE011/Binimetinib | 18 (14) | 93 (45-NA) |
| BYL719/Binimetinib | 43 (29) | 69 (42-82) |
| LEE011/Encorafenib | 33 (29) | 56 (34-131) |
| LEE011/Everolimus | 97 (58) | 51 (40-77) |
| BYL719/LEE011 | 39 (20) | 50 (39-NA) |
| Gemcitabine (50 mg/Kg) | 38 (31) | 44 (22-67) |
| LFW527/Everolimus | 33 (24) | 38 (34-83) |
| BKM120/Binimetinib | 66 (53) | 36 (27-44) |
| BKM120/LDE225 | 35 (28) | 35 (27-59) |
| BKM120/Encorafenib | 33 (28) | 33 (23-76) |
| Encorafenib/Binimetinib | 33 (31) | 33 (20-68) |
| Everolimus | 60 (51) | 33 (28-45) |
| BYL719/HSP990 | 64 (47) | 32 (26-51) |
| INC424/Binimetinib | 38 (31) | 32 (27-59) |
| LCL161/Paclitaxel | 29 (24) | 32 (23-48) |
| CKX620 | 70 (52) | 29 (21-39) |
| Binimetinib | 227 (183) | 28 (24-33) |
| BYL719/LJM716 | 203 (165) | 28 (26-31) |
| CLR457 | 212 (183) | 28 (25-33) |
| BKM120 | 224 (188) | 27 (24-28) |
| BYL719 | 211 (186) | 27 (24-32) |
| Trametinib | 38 (30) | 27 (21-43) |
| BYL719/Cetuximab | 40 (31) | 25.5 (19-43) |
| 5FU | 43 (36) | 24 (18-38) |
| BYL719/Cetuximab/Encorafenib | 42 (35) | 24 (15-39) |
| BYL719/Encorafenib | 41 (35) | 23 (16-38) |
| Cetuximab/Encorafenib | 42 (37) | 23 (11-36) |
| LFW527/Binimetinib | 71 (65) | 22 (17-29) |
| BKM120/LJC049 | 41 (30) | 21 (16-38) |
| Dacarbazine | 32 (23) | 21 (12-38) |
| LEE011 | 245 (217) | 21 (19-25) |
| Paclitaxel | 66 (60) | 20.5 (15-37) |
| BYL719/LGH447 | 29 (26) | 19 (14-29) |
| INC280/Trastuzumab | 43 (38) | 18 (14-21) |
| INC424 | 75 (75) | 18 (14-22) |
| LJM716/Trastuzumab | 83 (70) | 17 (14-22) |
| LLM871 | 110 (96) | 17 (14-21) |
| LJM716 | 101 (93) | 16 (14-21) |
| Cetuximab | 72 (68) | 15.5 (11-19) |
| Trastuzumab | 80 (76) | 15.5 (13-17) |
| Erlotinib | 29 (24) | 15 (8-35) |
| HDM201 | 191 (171) | 15 (14-17) |
| HSP990 | 92 (75) | 15 (14-20) |
| Tamoxifen | 39 (38) | 15 (14-17) |
| BGJ398 | 112 (104) | 14 (13-17) |
| CGM097 | 139 (128) | 14 (12-16) |
| Encorafenib | 76 (73) | 14 (11-19) |
| LDE225 | 33 (26) | 14 (11-16) |
| LFA102 | 39 (36) | 14 (13-20) |
| TAS266 | 33 (31) | 14 (9-22) |
| Figitumumab | 82 (78) | 13 (11-17) |
| LDK378 | 33 (32) | 13 (8-18) |
| LKA136 | 119 (115) | 13 (11-16) |
| WNT974 | 70 (61) | 12.5 (10-15) |
| Abraxane | 38 (38) | 12 (9-14) |
| INC280 | 71 (68) | 12 (10-14) |
| Binimetinib (3.5 mg/Kg) | 38 (36) | 11.5 (7-20) |
| Figitumumab/Binimetinib | 38 (35) | 11 (7-25) |
| LGW813 | 33 (32) | 11 (10-21) |
| LJC049 | 42 (42) | 11 (10-15) |
| LGH447 | 29 (29) | 10 (8-15) |

Of the 1103 trials we found that the biomarkers failed to capture the treatment effect in ~70% of them. A volcano plot showing how the study PFS HR relates to whether the treatment effect was captured or not can be seen in Figure 2. We can see that the likelihood of capturing the treatment effect does appear to depend on the size of the treatment effect, PFS HR. Although there is no guarantee that for a specific PFS HR range the biomarkers fully capture the treatment effect for all studies.

**Figure 2:**
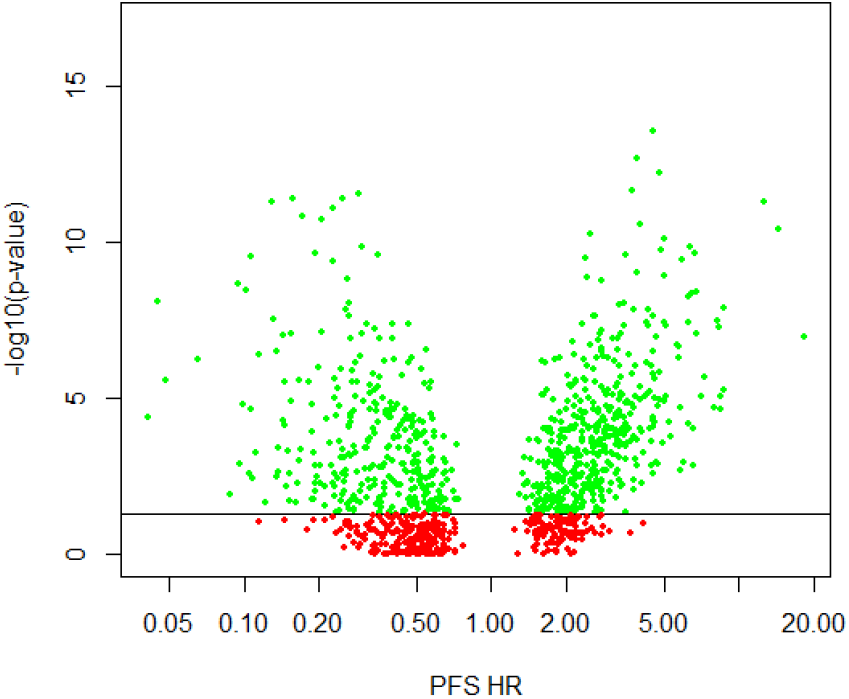
Plot showing the p-value and HR for each of the 1103 trials. Those in green had a p-value<0.05 and those in red did not.

## Discussion

The number of clinical analyses exploring surrogate markers is growing.^7–27^ These studies can be grouped into two categories those that explore the dynamics of tumour burden^12,25–27^ and those that explore other trial reported metrics such as PFS and Response Rate.^8,10,18,19^ Those based on the dynamics of tumour burden are predominantly based on one or two trials with only a handful exploring more^12^, whereas those based on current efficacy metrics are very large meta analyses.^8,10,18,19^ This paper is concerned with the studies exploring tumour dynamics as a surrogate biomarker. Most of those studies have never assessed whether the treatment effect was fully captured by the surrogate biomarker they proposed. Furthermore, the biomarkers being explored had never been assessed preclinically.

There are numerous advantages of exploring surrogate biomarkers in preclinical systems such as xenografts first before moving to patients. There is only one tumour to assess, which is clearly visible and easy to measure, unlike in a patient where quantifying initial level of tumour burden is very challenging. The measurement of the tumour can be done on a far more regular basis than the clinic as callipers are a far cheaper method than imaging scans. In general, the conditions are far more controlled preclinically than clinically. Therefore, if a biomarker does not perform well preclinically the chances of it performing well in more heterogenous and less controlled conditions like the clinic reduces dramatically. Indeed, the imaging biomarker community are aware of this and have developed a roadmap for new imaging biomarkers; a key part of this is performing a thorough preclinical evaluation before exploring a candidate biomarker in the clinic.

In this article we explored a large preclinical database of 61 drugs and 100s of diverse xenografts to explore how often the treatment effect is fully captured by potential surrogate biomarkers relating to tumour burden. The database extracted from the literature provided tumour burden time-series for xenografts for up to a year. Given these xenografts only had a single lesion the choice of biomarkers was limited. The current size and how the size had changed compared to the previous time-point were the only time-varying biomarkers considered. When these biomarkers were evaluated in this analysis only in 30% of the two-arm mouse trials did the biomarkers fully capture the treatment effect on disease progression times.

In summary, this study highlights how the emerging field of mouse clinical trial data-sets could be a testing ground for candidate surrogate biomarkers. An evaluation of one such database, the only one with open-access currently, presented herein has shown that surrogate biomarkers based purely on tumour burden measurements are unlikely to ever fully capture the treatment effect on the end-point. The glass is indeed less than half full.

## References

1. Driscoll, J.J. & Rixe, O. Overall survival: still the gold standard: why overall survival remains the definitive end point in cancer clinical trials. Cancer J 15, 401–405 (2009).

2. Prentice, R. L. Surrogate endpoints in clinical trials: definition and operational criteria. Stat Med 8, 431–440 (1989).

3. Dowden, H. & Munro, J. Trends in clinical success rates and therapeutic focus. Nature Reviews Drug Discovery 18, 495–496 (2019).

4. Gao, H. et al. High-throughput screening using patient-derived tumor xenografts to predict clinical trial drug response. Nat. Med. 21, 1318–1325 (2015).

5. Migliardi, G. et al. Inhibition of MEK and PI3K/mTOR Suppresses Tumor Growth but Does Not Cause Tumor Regression in Patient-Derived Xenografts of RAS-Mutant Colorectal Carcinomas. Clin Cancer Res 18, 2515–2525 (2012).

6. Schwartz, L. H. et al. RECIST 1.1 – Update and Clarification: From the RECIST Committee. Eur J Cancer 62, 132–137 (2016).

7. Adam, R. Surrogate Markers of Overall Survival in Metastatic Colorectal Cancer: An Evolving Challenge Still More Complex with Repeat Surgery. Ann Surg Oncol 21, 1763–1764 (2014).

8. Anand, S. et al. A systematic review of surrogate endpoints (SEPs) for overall survival (OS) in metastatic colorectal cancer (mCRC). JCO 37, e18206–e18206 (2019).

9. Berghmans, T. et al. Surrogate markers predicting overall survival for lung cancer: ELCWP recommendations. European Respiratory Journal 39, 9–28 (2012).

10. Blumenthal, G. M. et al. Overall Response Rate, Progression-Free Survival, and Overall Survival With Targeted and Standard Therapies in Advanced Non-Small-Cell Lung Cancer: US Food and Drug Administration Trial-Level and Patient-Level Analyses. JCO 33, 1008–1014 (2015).

11. Brady-Nicholls, R. et al. Prostate-specific antigen dynamics predict individual responses to intermittent androgen deprivation. Nature Communications 11, 1–13 (2020).

12. Claret, L., Mercier, F., Houk, B. E., Milligan, P. A. & Bruno, R. Modeling and simulations relating overall survival to tumor growth inhibition in renal cell carcinoma patients. Cancer Chemother Pharmacol 76, 567–573 (2015).

13. Claret, L. et al. Model-based prediction of progression-free survival in patients with first-line renal cell carcinoma using week 8 tumor size change from baseline. Cancer Chemother. Pharmacol. 78, 605–610 (2016).

14. Desmée, S., Mentré, F., Veyrat-Follet, C., Sébastien, B. & Guedj, J. Nonlinear joint models for individual dynamic prediction of risk of death using Hamiltonian Monte Carlo: application to metastatic prostate cancer. BMC Med Res Methodol 17, 1–12 (2017).

15. Heller, G. et al. Circulating Tumor Cell Number as a Response Measure of Prolonged Survival for Metastatic Castration-Resistant Prostate Cancer: A Comparison With Prostate-Specific Antigen Across Five Randomized Phase III Clinical Trials. Journal of Clinical Oncology (2017) doi:10.1200/JCO.2017.75.2998.

16. Litière, S. et al. The components of progression as explanatory variables for overall survival in the Response Evaluation Criteria in Solid Tumours 1.1 database. European Journal of Cancer 50, 1847–1853 (2014).

17. Mistry, H. B. On the relationship between tumour growth rate and survival in non-small cell lung cancer. PeerJ 5, e4111 (2017).

18. Pasalic, D. et al. Progression-free survival is a suboptimal predictor for overall survival among metastatic solid tumour clinical trials. European Journal of Cancer 136, 176–185 (2020).

19. Prasad, V., Kim, C., Burotto, M. & Vandross, A. The Strength of Association Between Surrogate End Points and Survival in Oncology: A Systematic Review of Trial-Level Meta-analyses. JAMA Intern Med 175, 1389–1398 (2015).

20. Randall, L. M. et al. Predictive value of serum CA-125 levels in patients with persistent or recurrent epithelial ovarian cancer or peritoneal cancer treated with bevacizumab on a Gynecologic Oncology Group phase II trial. Gynecol Oncol 124, 563–568 (2012).

21. Scher, H. I. et al. Circulating tumor cell biomarker panel as an individual-level surrogate for survival in metastatic castration-resistant prostate cancer. J. Clin. Oncol. 33, 1348–1355 (2015).

22. Sostelly, A. & Mercier, F. Tumor Size and Overall Survival in Patients With Platinum-Resistant Ovarian Cancer Treated With Chemotherapy and Bevacizumab. Clin Med Insights Oncol 13, (2019).

23. Stein, W. D. et al. Tumor Regression and Growth Rates Determined in Five Intramural NCI Prostate Cancer Trials: The Growth Rate Constant as an Indicator of Therapeutic Efficacy. Clin Cancer Res 17, 907–917 (2011).

24. Verbel, D. A., Heller, G., Kelly, W. K. & Scher, H. I. Quantifying the Amount of Variation in Survival Explained by Prostate-specific Antigen. Clin Cancer Res 8, 2576–2579 (2002).

25. Yu, M., Taylor, J. M. G. & Sandler, H. M. Individual Prediction in Prostate Cancer Studies Using a Joint Longitudinal Survival-Cure Model. Journal of the American Statistical Association 103, 178–187 (2008).

26. Zhang, J. et al. Relationship Between Tumor Size and Survival in Non–Small-Cell Lung Cancer (NSCLC): An Analysis of the Surveillance, Epidemiology, and End Results (SEER) Registry. Journal of Thoracic Oncology 10, 682–690 (2015).

27. Zheng, Y. et al. Population Modeling of Tumor Kinetics and Overall Survival to Identify Prognostic and Predictive Biomarkers of Efficacy for Durvalumab in Patients With Urothelial Carcinoma. Clinical Pharmacology & Therapeutics 103, 643–652 (2018).

